# T Residues Preceded by Runs of G are Hotspots of T→G Mutation in Bacteria

**DOI:** 10.1101/2022.10.28.514265

**Authors:** Joshua L. Cherry

## Abstract

The rate of mutation varies among positions in a genome. Local sequence context can affect the rate, and has different effects on different types of mutation. Here I report an effect of local context that operates to some extent in all bacteria examined: the rate of T→G mutation is greatly increased by preceding runs of three or more G residues. The strength of the effect increases with the length of the run. In *Salmonella*, in which the effect is strongest, a G run of length three increases the rate by a factor of ~26, a run of length four increases it by almost a factor of 100, and runs of length five or more increase it by a factor of more than 400 on average. The effect is much stronger when the T is on the leading rather than the lagging strand of DNA replication. Several observations eliminate the possibility that this effect is an artifact of sequencing error.

## Introduction

The rate of mutation varies among nucleotide sites within a genome. In addition to moderate variation among the bulk of sites, some sites, often referred to as hotspots, exhibit very high mutation rates. Often only mutation to a particular nucleotide is accelerated at a hotspot.

Some hotspots of mutation are caused by DNA modification. Methylation of cytosine at the C5 position increases the rate of C→T mutation in bacteria (Coulondre et al. 1978; Cherry 2018) and eukaryotes (Bird 1980; Cooper and Youssoufian 1988). N6-methylated adenines in Dam sites also exhibit an elevated mutation rate (Lee et al. 2012). Certain N4-methylated cytosines are hotspots of mutation, though the causal role of this methylation is uncertain (Cherry 2021).

Repeats of short sequence motifs, including homopolymer runs, are subject to high rates of insertion and deletion of one or more repeat units. The mechanism is thought to involve slipped-strand mispairing, in which the strands of the repeat region pair out of register near the end of the strand being synthesized (Levinson and Gutman 1987).

Although natural selection may generally favor lower mutation rates, it may favor hotspots at certain locations, particularly if reversal is facile or the mutation rate is high only in somatic cells. Bacterial “contingency loci” in which insertion and deletion occur frequently in homopolymer runs (Moxon et al. 2006) are examples. Reversibility need not be an issue if the benefits of hypermutation are due mainly to kin selection, i.e., they mostly accrue not to the mutant and its descendants, but to nonmutant close relatives (Cherry 2020).

Mutation rate and its determinants have been studied using natural variation and laboratory experiments. Natural variation, even within a species, is subject to confounding effects of selection if long evolutionary distances are involved, but it reflects mutational processes as they occur in nature. Laboratory experiments include mutation accumulation experiments, in which the effects of selection are minimized; laboratory evolution experiments, in which the main goal is adaptation; and selection experiments in which only mutants of a particular type survive. Experiments with mutator genotypes provide clues to mechanisms of mutation and heterogeneity of its rate.

Studies of various types have revealed effects of nearest-neighbor nucleotides (the bases immediately preceding and following the mutated position) on mutation rate (Lee et al. 2012; Sung et al. 2015; Goncearenco et al. 2017). The identities of flanking bases have different effects on different types of mutations, e.g., C→T vs. C→A and C→G.

Factors other than local sequence context have been found to correlate with mutation rate. These include orientation with respect to the direction of replication (leading/lagging strand asymmetry) (Lujan et al. 2014; Sung et al. 2015), the rate and direction of transcription (Hudson et al. 2003; Lujan et al. 2014), distance from the origin of replication (Mira and Ochman 2002; Lujan et al. 2014), and nucleosome positioning in eukaryotes (Lujan et al. 2014; Li and Luscombe 2020).

Here I describe a mutational phenomenon that occurs in bacteria: the rate of T→G mutation is greatly increased if the T is immediately preceded by three or more G resides. This effect was detected in all bacteria analyzed. It was inferred mainly from sequence comparisons between closely related natural isolates, which mitigates the confounding effects of selection. In *Salmonella*, in which the effect is strongest, this phenomenon accounts for about one third of all T→G mutations.

## Results

### Elevated T→G Mutation Rate at GGGT Sites in *Salmonella*

Examination of candidate hotspots of mutation in *Salmonella* revealed that many exhibit mainly T→G changes at GGGT. This observation motivated a formal analysis of the rate of changes of this type.

Analysis of *Salmonella* data from the NCBI Pathogens project (PDG000000002.2479) yielded 1,494,207 polarizable single-nucleotide changes meeting criteria for strength of support. Of the 102,153 T→G changes, 36,130, or ~35%, were at positions with at least three immediately preceding G residues, though these constitute only 1.1% of the T positions.

Table 1 presents relative T→G rates for positions preceded by G tracts of different lengths. The rate is increased substantially by G tracts of length three, and even more by longer tracts. Because T→G transversions constitute only ~21% of point mutations at T positions not preceded by G, the factor by which the total mutation rate is increased is about five-fold lower, but nevertheless quite large.

**Table 1.** Effects of preceding G tracts on T->G mutation rate in *Salmonella*

|  | Length of Preceding G Tract |  |  |  |  |
| --- | --- | --- | --- | --- | --- |
|  | 1 | 2 | 3 | 4 | 5 |
| Mutations | 6953 | 3439 | 16527 | 9920 | 9683 |
| Sites | 367102.6 | 120477.2 | 20557.5 | 3287.9 | 733.2 |
| Relative rate | 0.62 | 0.93 | 26.3 | 98.6 | 431 |
| 95% CI | 0.60-0.63 | 0.90-0.97 | 25.8-26.7 | 96.5-101 | 422-441 |

### Evidence Against Systematic Sequencing Error

Systematic sequencing error might produce the appearance of high mutation rates at certain sequence motifs. Several types of analysis were performed to test for this possibility.

A mutation may occur along an internal branch of the phylogenetic tree, usually affecting multiple isolates that constitute a clade. In contrast, a sequencing error at a particular position is unlikely to affect multiple closely related isolates. The fraction of mutations of a particular type inferred to have occurred along internal branches is therefore an indication of whether they are genuine. This fraction is 32.7% for mutations other than those of interest. For mutations of interest this fraction is very similar: 32.7%, 33.3%, and 32.1% for T→G mutations at sites preceded by G runs of length three, four, and five or more. This result suggests that the mutations of interest are mostly genuine.

A related analysis considers only differences between pairs of relatively distant isolates from the same cluster. For each cluster with at least 2000 inferred mutations, two isolates, one descended from each child node of the root of the tree, were chosen at random, and only sequences differences between these pairs were considered. This procedure reduces the ratio of genome sequences to inferred mutations by more than a factor of five. Because the total number of sequencing errors is expected to be proportional to the number of genome sequences, the fraction of sequencing errors among inferred mutations should be reduced by a similar factor. Nevertheless, the fraction of mutations of interest is as large as in the full dataset: 2.48% on average among 10,000 random choices of isolate pairs, compared to 2.42% for the full dataset. Of these, 26.5% are at positions with four preceding G residues, and 26.3% at positions with five or more, similar to the fractions in the full dataset, 27.5% and 26.8%.

The number of mutations of interest inferred from these isolate pairs should correlate with the number of other mutations if mutations of both types are mostly genuine, since both will tend to increase with the evolutionary time separating the isolates. This correlation will be imperfect due to stochasticity of mutation. It can be compared to the correlation for counts simulated under the assumption that the expected numbers correlate perfectly (see Methods). The average Spearman correlation for the actual counts is 0.265, remarkably close to the average for simulated counts, 0.263. The expected numbers appear to correlate nearly perfectly, as expected for genuine mutations, but not for sequencing errors.

Although motifs including runs of G are a source of sequence-specific errors for some Illumina platforms (Stoler and Nekrutenko 2021), only reads in one direction are affected. Because the analysis in Table 1 considered only mutations with strong bidirectional support (at least 20 aligned reads, at least 90% of which supported the called base, and at least 25% of them supporting reads in each direction), such errors should mostly be eliminated. Furthermore, removing all requirements for read support does not decrease the apparent strength of the effect, instead reducing it slightly (results not shown). Making the requirements more stringent (at least 50 reads, 100% supporting, and at least 40% in each direction) also does not change the strength of the effect greatly.

The frequencies of different types of errors vary among sequencing technologies. Illumina NovaSeq, unlike other Illumina platforms, does not exhibit elevated error rates after homopolymer runs (Stoler and Nekrutenko 2021). PacBio sequencing is relatively free of systematic non-indel errors (Weirather et al. 2017) and usually does not involve an amplification step, a potential source of errors. Although the technology is incompletely specified in the metadata for most of the genomes, some determined solely by one of these technologies can be identified. A total of 1132 mutations were supported by at least one genome for which NovaSeq is listed as the only sequencing technology. Of these, 30, or 2.65%, were of the type of interest, close to the 2.42% found among all mutations. Of these, eight were at positions with four preceding G residues, and four at positions with five or more, consistent within sampling variance with the distribution in Table 1. Of the 5148 mutations supported by at least one genome determined solely by PacBio sequencing, 118, or 2.29% (95% CI 1.90-2.74%), were of the type of interest, similar to the fraction in the full set. Of these, 26 were at positions with four preceding G residues, and 39 at positions with five, again in accord with Table 1. All of these results support a genuine phenomenon rather than sequencing error.

### Confirmation with Other Datasets

The genomes of hundreds of *E. coli* isolates from a laboratory evolution experiment have been sequenced, and changes from the ancestral sequence reported (Tenaillon et al. 2016). The total number of inferred sequence changes is relatively small. In addition, strong selection and possible differences in mutational spectrum between laboratory and natural growth may affect the results. However, the availability of multiple genome sequences for each replicate, and of sequences from different time points, allows additional tests that strongly discriminate between genuine changes and sequencing errors.

Table 2 presents relative rate estimates for mutations of the type of interest in non-mutator isolates. Despite the small absolute numbers (only 21 GGGT->GGGG changes), a strong effect of preceding G runs on T->G mutation rate is evident.

**Table 2.**
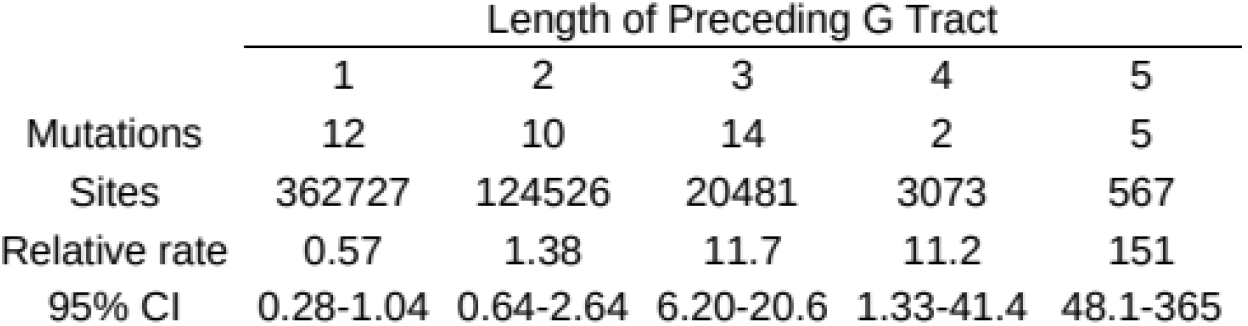
Effects of preceding G tracts on T->G mutation rate in the *E. coli* evolution experiment

|  | Length of Preceding G Tract |  |  |  |  |
| --- | --- | --- | --- | --- | --- |
|  | 1 | 2 | 3 | 4 | 5 |
| Mutations | 12 | 10 | 14 | 2 | 5 |
| Sites | 362727 | 124526 | 20481 | 3073 | 567 |
| Relative rate | 0.57 | 1.38 | 11.7 | 11.2 | 151 |
| 95% CI | 0.28-1.04 | 0.64-2.64 | 6.20-20.6 | 1.33-41.4 | 48.1-365 |

The detection of a sequence change in more than one isolate from a single replicate is evidence that it is genuine. Of the 21 differences of interest, 13, or 62%, were found in more than one isolate from the replicate, higher than the fraction among other sequence differences, 55%. In contrast, only two changes of this type were found in genomes from different replicates, despite the fact that there are many more between-than within-replicate pairs. Both of the changes with a G run of length four, and one of the five with a run of length five or more, were found in more than one genome sequence within the replicate.

Genuine sequence differences, but not sequencing errors, are expected to increase in number with the progress of the evolution experiment. Fig. 1 shows how the frequency of GGGT→GGGG differences from the ancestor changes with time. Their frequency is zero at the three earliest time points and increases nearly monotonically thereafter.

**Fig. 1.**
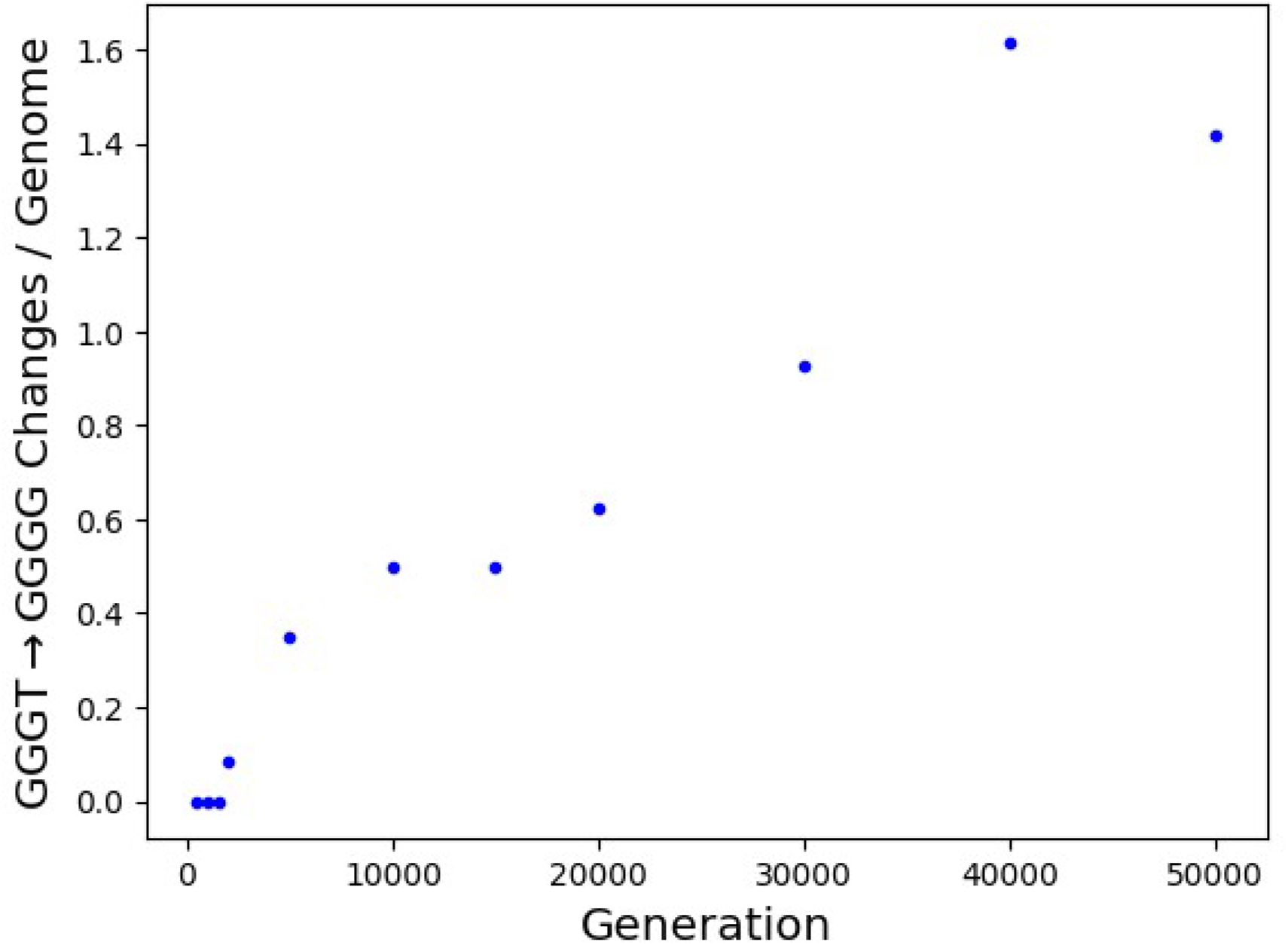
GGGT →GGGG mutation frequency in the *E. coli* long-term evolution experiment as a function of the number of generations of evolution. Division by the number of genomes accounts for the exclusion of isolates with mutator mutations.

The phenomenon is also apparent in the mutation accumulation experiment reported in the same work. Of the 197 single-base mutations detected, 13 are T→G transversions. Three of these are at positions preceded by three or more G residues. GGGT →GGGG mutations therefore constitute 24% of T→G mutations and 1.5% of all single-base mutations, in rough agreement with the results for natural isolates of *Salmonella*. The numbers of preceding G residues are three, four, and six.

Marvig et al. (2015) sequenced the genomes of *Pseudomonas aeruginosa* isolated serially from cystic fibrosis patients. As shown in Table 3, the observed rate of T→G change is elevated by preceding G runs. The changes of interest are found in multiple genome sequences from the same clone type (restricted to one or a few patients) in 66.7% of cases, compared to 53.5% for other types of single-base changes. Changes of interest are found in multiple clone types at only three positions, so this is strong evidence that the changes are genuine.

**Table 3.** Effects of preceding G tracts on T->G mutation rate in *P. aeruginosa* serial isolates

|  | Length of Preceding G Tract |  |  |  |  |
| --- | --- | --- | --- | --- | --- |
|  | 1 | 2 | 3 | 4 | 5 |
| Mutations | 23 | 16 | 33 | 14 | 10 |
| Sites | 365391 | 187607 | 39530 | 7718 | 1304 |
| Relative rate | 0.88 | 1.19 | 11.7 | 25.3 | 107 |
| 95% CI | 0.53-1.39 | 0.66-2.02 | 7.64-17.4 | 13.4-44.4 | 49.9-205 |

If the apparent changes of interest are mostly genuine, the number observed in an isolate should correlate with the number of point mutations of other types, which varies considerably among isolates. Because isolates from the same clone type are not independent, one non-mutator isolate from each clone type was chosen at random for computation of this correlation. This procedure was repeated 100,000 times. The Spearman correlation was on average 0.631. This is very close to 0.637, the average for counts simulated under the assumption that their expected values are perfectly correlated. These results indicate a near perfect correlation between the expected counts of mutations of interest and of other mutations, which is strong evidence that almost all of the mutations of interest are real.

### Leading/Lagging Strand Asymmetry

Replication of a circular bacterial chromosome generally proceeds in both directions from an origin of replication, terminating at approximately the opposite point of the circle. The replication fork is asymmetric, and the mutational process differs between leading and lagging strand synthesis (Fijalkowska et al. 1998). I therefore assessed the T→G mutation rate in *Salmonella* as a function of orientation with respect to the direction of replication.

Many of the genome assemblies, including the reference assemblies for some clusters, are highly fragmented. Along with the fact that many genome rearrangements have occurred within *Salmonella*, this makes it impossible to know the orientation of SNP positions in some clusters. I therefore made use of the *Salmonella* cluster with the largest number of inferred mutations, PDS000089910.229, which contains 10,062 isolates and yields 66,534 chromosomal mutations for analysis. The use of a single cluster also makes it easier to know that different changes have occurred at the same position in the genome.

Fig. 2, top panel, shows GT skew for the sequence of the reference chromosome for the cluster (CP093400.1), along with the approximate locations of the origin and terminus of replication determined from its minimum and maximum. The location of the origin is approximately consistent with the location of the *dnaA* gene at the beginning of the sequence and its forward-strand orientation, as the origin is generally not far upstream of this gene in *Salmonella*.

**Fig. 2.**
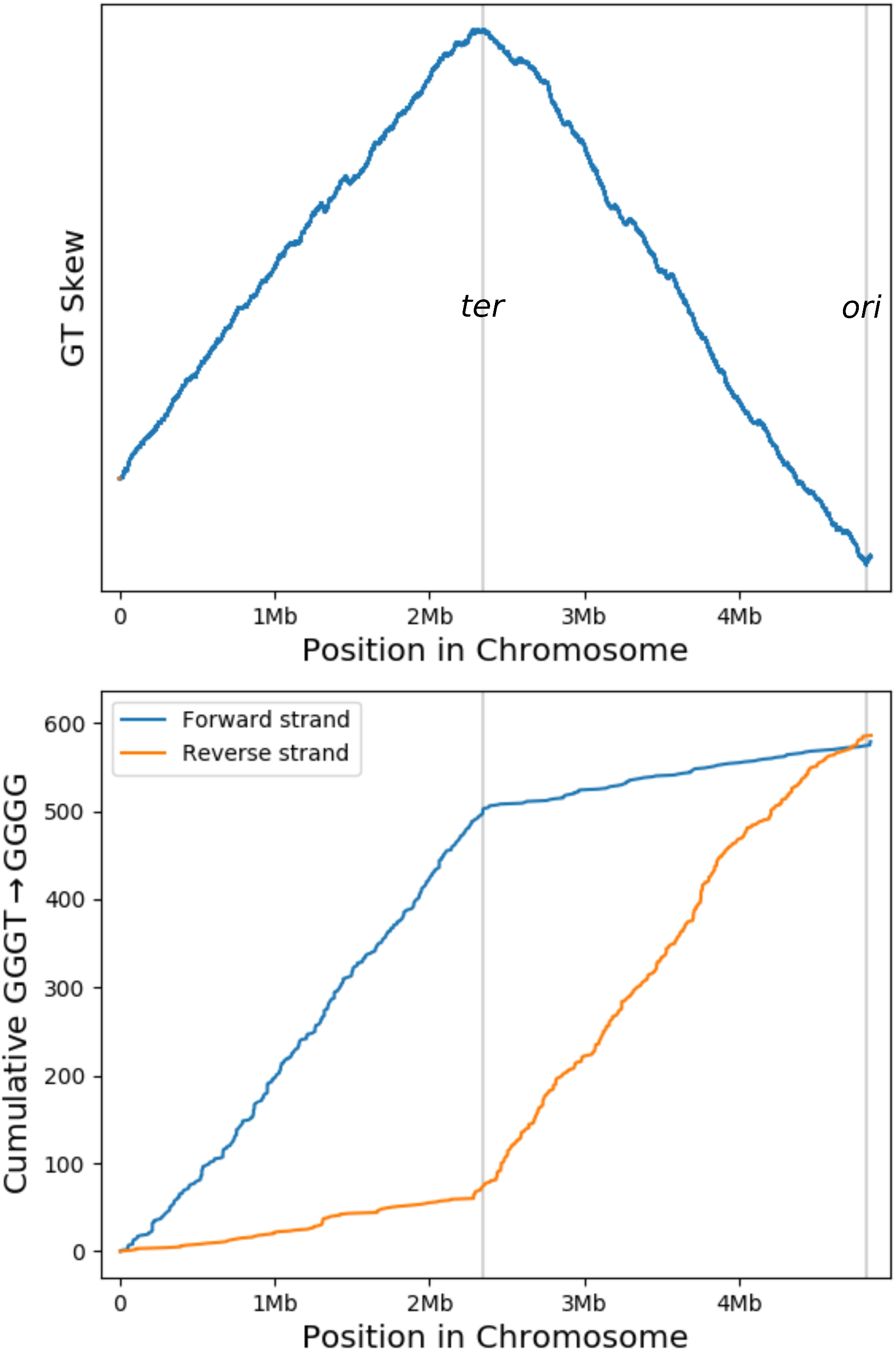
Orientation-dependence of GGGT →GGGG mutation in *Salmonella*. The upper panel shows GT skew, which reveals the approximate locations of the origin and terminus of replication. The lower panel shows the cumulative number of mutations of interest for the forward and reverse orientations.

The lower panel of fig. 2 shows the cumulative distribution of the genome position of T→G changes at positions preceded by three or more G residues for “forward” and “reverse” orientations. Also shown are the approximate positions of the origin and terminus of replication. It is evident that there are many more changes at sites where GGGT rather than its complement is on the leading strand. Table 4 shows that the rate per site after G runs of length three or more is much higher when the T is on the leading strand.

**Table 4.** Orientation-dependence of T->G mutation rate in *Salmonella*

|  |  | Length of Preceding G Tract |  |  |  |  |  |
| --- | --- | --- | --- | --- | --- | --- | --- |
|  |  | 0 | 1 | 2 | 3 | 4 | 5 |
| Leading strand | Mutations | 1343 | 183 | 133 | 434 | 299 | 287 |
|  | Sites | 899234 | 190804 | 64524 | 10876 | 1928 | 486 |
|  | Relative rate | 1.08 | 0.69 | 1.49 | 28.9 | 112 | 428 |
|  | 95% CI | 1.01-1.16 | 0.59-0.81 | 1.24-1.78 | 26.1-32.0 | 99.3-127 | 377-484 |
| Lagging strand | Mutations | 1148 | 155 | 63 | 99 | 27 | 21 |
|  | Sites | 905655 | 175566 | 55669 | 9804 | 1374 | 247 |
|  | Relative rate | 0.92 | 0.64 | 0.82 | 7.32 | 14.2 | 61.6 |
|  | 95% CI | 0.86-0.99 | 0.54-0.75 | 0.63-1.05 | 5.92-8.95 | 9.36-20.8 | 38.1-94.4 |
| Rate ratio<sup>a</sup> |  | 1.18 | 1.09 | 1.82 | 3.95 | 7.89 | 6.95 |
| 95% CI |  | 1.09-1.28 | 0.87-1.35 | 1.34-2.50 | 3.17-4.97 | 5.32-12.2 | 4.46-11.4 |

<sup>a</sup>Ratio of rate per site on leading and lagging strand

### Effects of G Runs on Other Types of Mutation

Fig. 3 shows relative rates of other categories of mutation at positions adjacent to G tracts of different lengths in *Salmonella*. Values are relative to the rate with a G tract of length zero, that is, for positions preceded or followed by a base other than G.

**Fig. 3.**
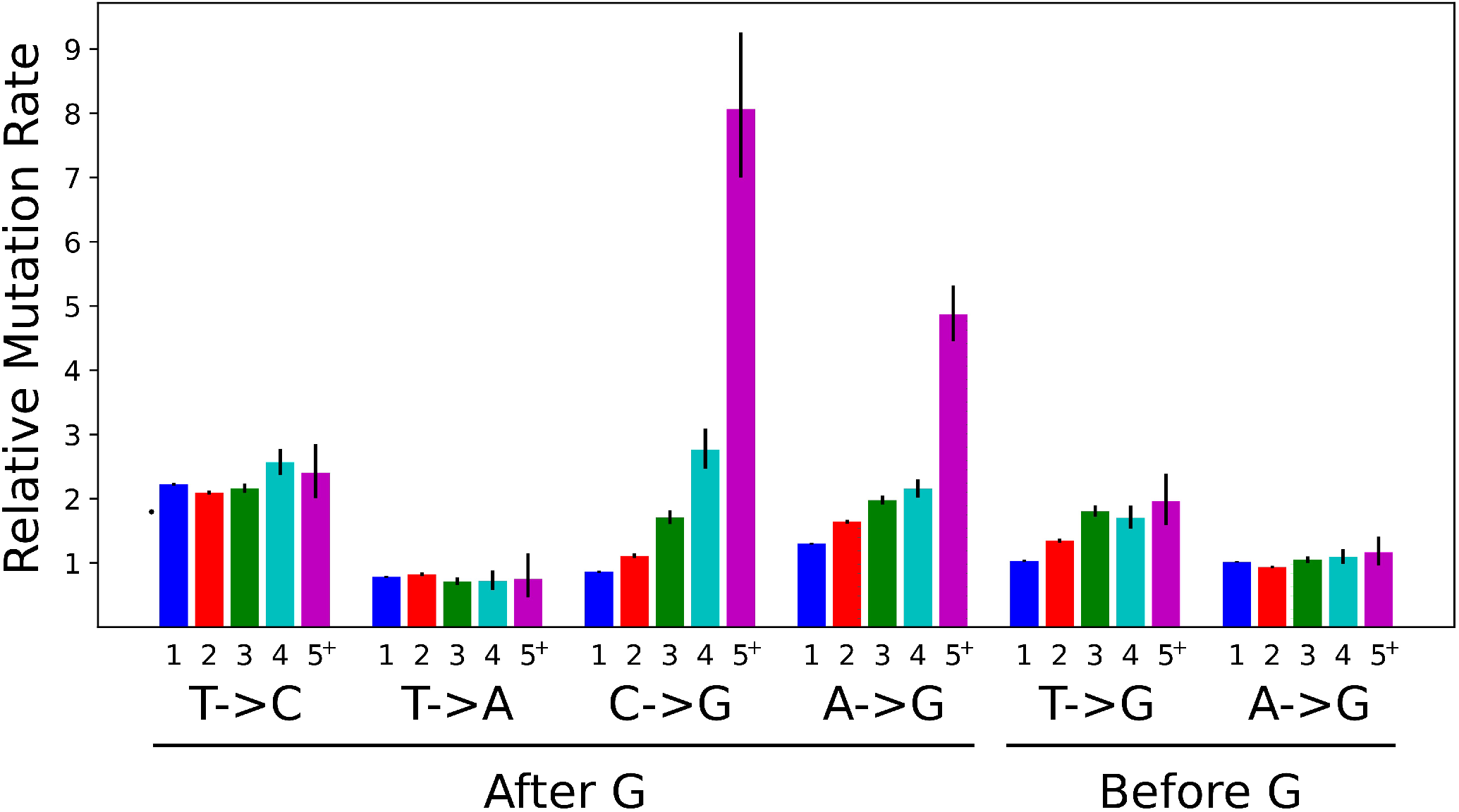
Relative rates of various types of mutation at positions adjacent to G tracts of different lengths in *Salmonella*. Rates are relative to positions not adjacent to G. Error bars indicate 95% confidence intervals. “5^+^” denotes five or more G residues.

Mutation of T to C or A is not strongly affected by preceding runs of G, i.e., their effect is specific for mutation to G. Although the rate of C→G mutation is about twofold higher when the preceding base is a G, it does not increase substantially with additional preceding G residues.

An effect of mutation of C or A to G is evident, particularly for longer runs, but it is weak compared to the effect on mutation of T. Although the effect for runs of length four or more appears to be stronger for C than for A, this is attributable to the larger denominator for A, which reflects the generally higher rate of transitions compared to transversions. In terms of absolute numbers of additional mutations per site, the effect on A is larger than that on C, but still much smaller than the effect on T. The apparent strength for runs of at least five may also be affected by differences between preceding run length distributions among nucleotides.

G runs that follow rather than precede a T have only a small effect on T→G mutation rate. There is no substantial effect of following G runs on A→G mutation. Equivalently, preceding runs of C do not substantially affect the rate of T→C mutation.

### Contribution to Nearest Neighbor Effects

Fig. 4 shows the effects of nearest neighbors (the immediately preceding and following nucleotides) on rates of different types of mutation in *Salmonella*. All mutations are oriented so that the ancestral base is a pyrimidine. The contribution of GGGT →GGGG mutations is shown in red. Also shown are the contributions of C→T mutation at Dcm methylation sites and those of mutation at Dam methylation sites (represented as mutation of the paired T).

**Fig. 4.**
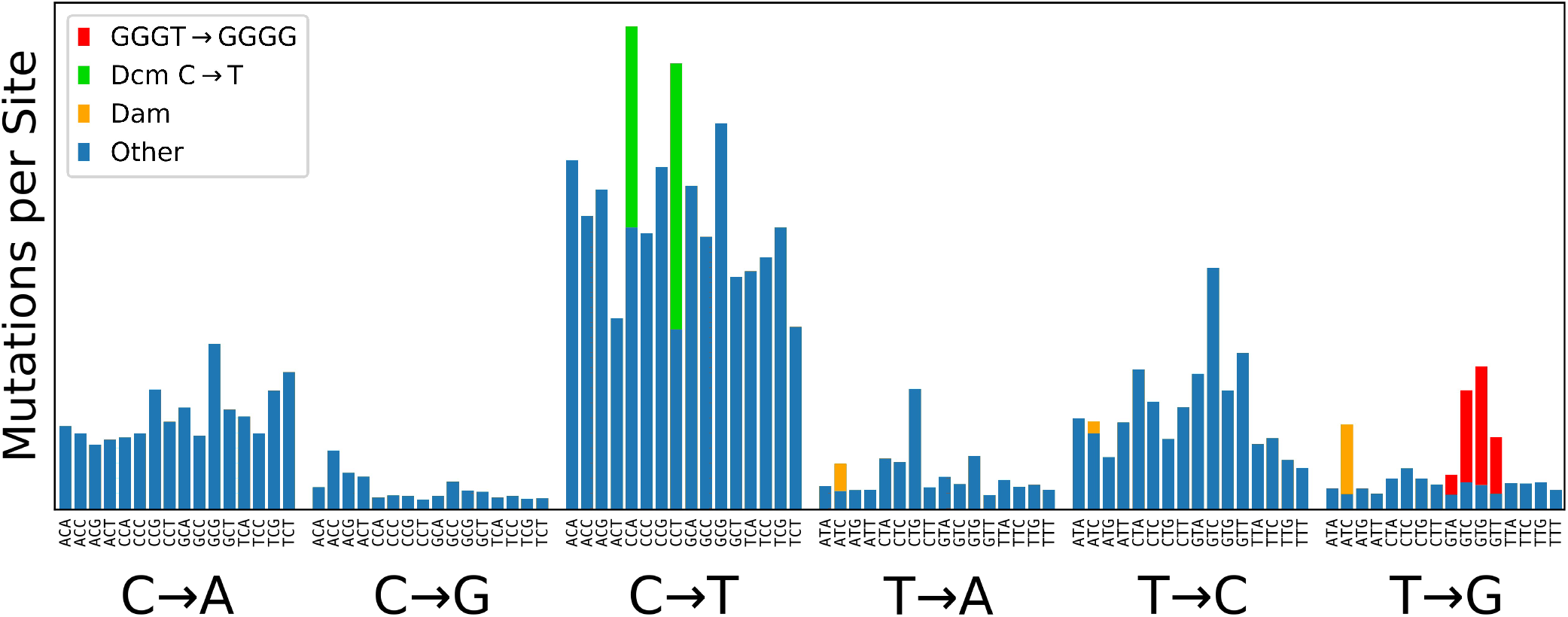
Context-dependence of mutation rates in *Salmonella*. For each strand-symmetrized category of point mutation, the number of mutations per site is shown for all sixteen combinations of preceding and following base. The contributions of GGGT->GGGG mutations, transitions at Dcm-methylated sites, and mutations opposite Dam-methylated adenines are indicated by color.

### Other Bacteria

Other bacteria were assessed for an effect of preceding G runs on the rate of T→G mutation using the NCBI Pathogens data. The choice of taxa for analysis was guided by the quantities of available data, the desire for phylogenetic diversity, and the desire to analyze close relatives of *Salmonella*.

Fig. 5 presents results for several bacteria. The phenomenon operates to some extent in all of these, though its strength varies considerably among them (note that the vertical axis is logarithmic). The effect is strongest in *Salmonella*, followed by its close relative *E. coli*.

**Fig. 5.**
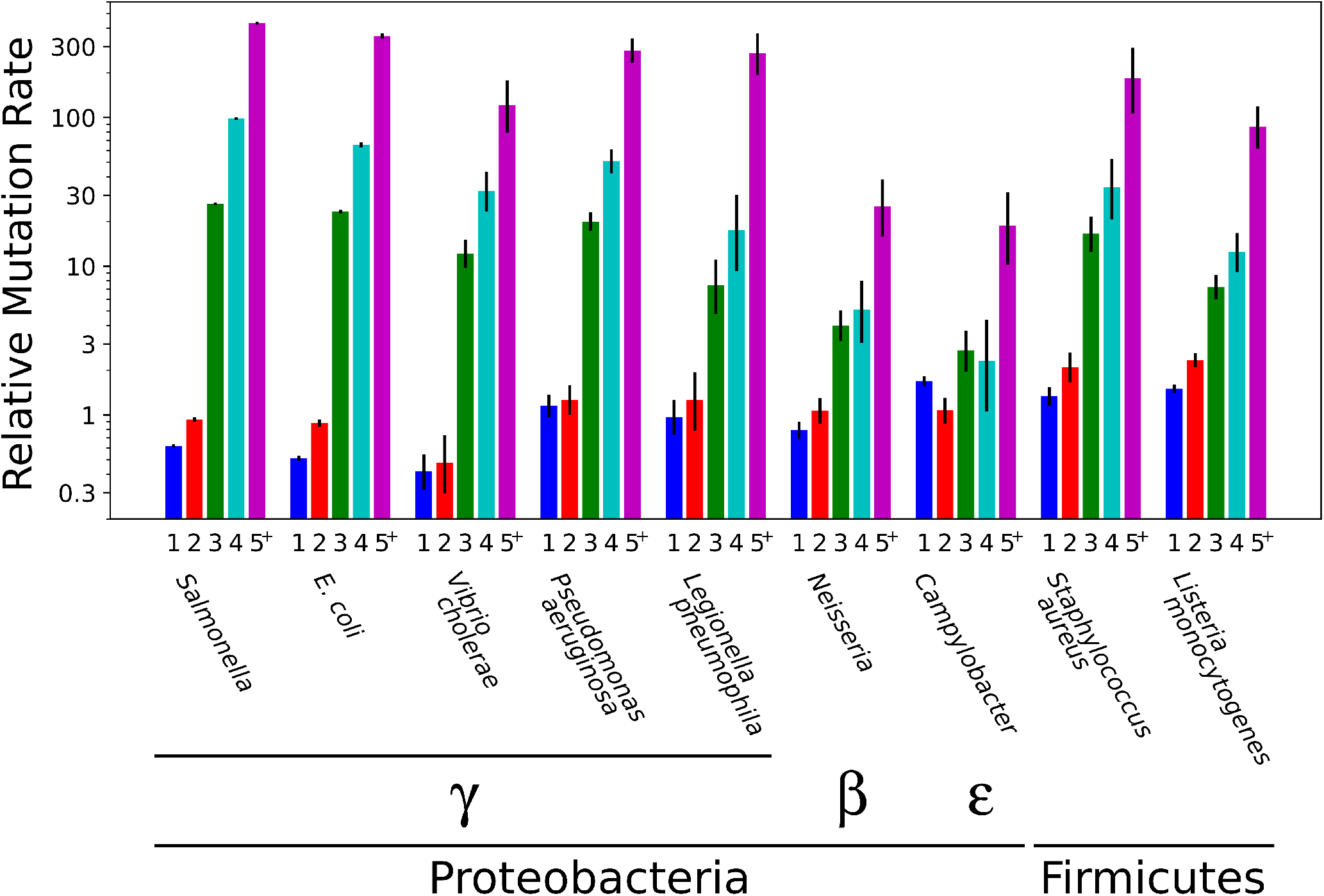
Relative T->G mutation rate with different numbers of preceding G residues in various bacteria. Values are relative to the rate with no preceding G. Error bars indicate 95% confidence intervals. “5^+^” denotes five or more G residues. Note the logarithmic vertical scale.

### No Strong Effect in Yeast

An analysis of data from a yeast mutation accumulation experiment (Sharp et al. 2018) is shown in Table 5. No GGGT → GGGG mutations were observed. This result is consistent with no enhancement of transversion rate by preceding runs of G. Nonetheless, a sizable effect of four or more G residues cannot be ruled out due to the low absolute numbers of other types of mutations. For just three, however, the 95% confidence interval limits the size of any enhancement to about a factor of three. If all sites with G runs of length three or more are combined, a tighter upper bound of 2.65 is obtained. Assuming that additional G residues do not decrease the mutation rate, this bound can be applied to runs of length three. The implication is that any effect of three preceding G residues in yeast is weaker than that in any of the bacteria analyzed, with the possible exception *Campylobacter*. Use of single-sided confidence intervals, arguably justified for counts of zero, would reduce all of the upper bounds by ~20%, strengthening this conclusion.

**Table 5.**
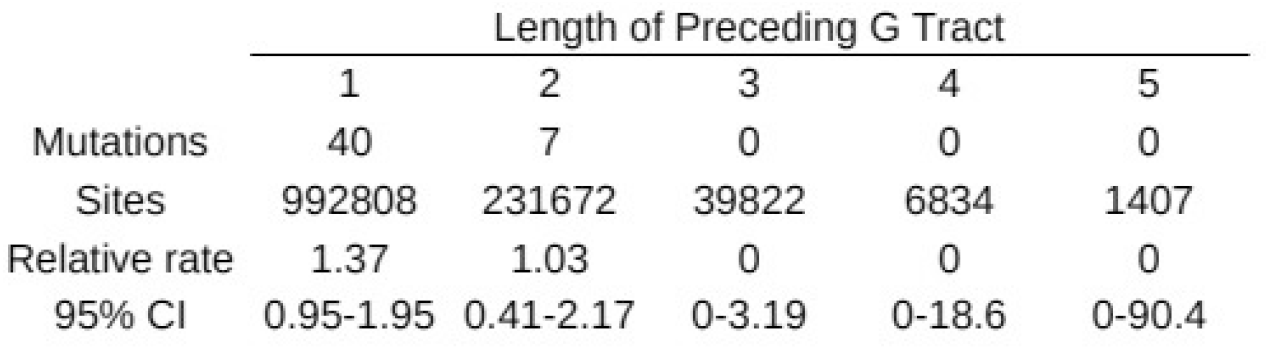
Effects of preceding G tracts on T->G mutation rate in yeast

### Effect of MutT Deficiency

The MutT protein catalyzes the hydrolysis of 8-oxo-dGTP, reducing the rate of T→G mutation that results from misincorporation of 8-oxo-G across from A in the template (Setoyama et al. 2011). This pathway is responsible for most mutations in strains deficient in *mutT*. Couce et al. (2017) compared the mutational spectra of *E. coli* non-mutator and mutator strains, including *mutT* mutants.

As demonstrated above, the effect of G runs on T→G mutation is apparent during laboratory growth of non-mutator *E. coli*. Table 6 show analogous results for *mutT*-deficient strains in mutation accumulation and long-term evolution experiments. Mutation rates are largely unaffected by preceding runs of G in *mutT* mutants, strongly suggesting that G runs do not exert their effect through incorporation of 8-oxo-G.

**Table 6.** Effects of preceding G tracts on T->G mutation rate in *mutT* mutants of *E. coli*

|  |  | Length of Preceding G Tract |  |  |  |  |
| --- | --- | --- | --- | --- | --- | --- |
|  |  | 1 | 2 | 3 | 4 | 5 |
| Mutation Accumulation | Mutations | 144 | 67 | 15 | 4 | 2 |
|  | Sites | 363072 | 124766 | 20749 | 3102 | 581 |
|  | Relative rate | 0.35 | 0.48 | 0.64 | 1.15 | 3.06 |
|  | 95% CI | 0.30-0.42 | 0.37-0.61 | 0.36-1.06 | 0.31-2.94 | 0.37-11.1 |
| Long-term Evolution | Mutations | 387 | 218 | 33 | 6 | 3 |
|  | Sites | 362727 | 124526 | 20481 | 3073 | 567 |
|  | Relative rate | 0.33 | 0.55 | 0.50 | 0.61 | 1.65 |
|  | 95% CI | 0.30-0.37 | 0.48-0.63 | 0.35-0.71 | 0.22-1.33 | 0.34-4.84 |

## Discussion

The rate of T→G mutation in bacteria is especially high at positions immediately preceded by at least three G residues and increases with the number of these. This phenomenon was apparent in all of the bacteria analyzed, which are phylogenetically diverse, but its strength varies considerably among them. It is strongest in *Salmonella*, and nearly as strong in the closely related *E. coli*. In *Salmonella*, the T→G rate at such sites is four-to eight-fold higher when the T, rather than the paired A, is on the leading strand of DNA replication.

Several types of evidence indicate that the phenomenon is not an artifact of systematic sequencing error. These include the high frequencies with which mutations of the type of interest are detected in two or more appropriately related isolates, increases in their numbers with time and with sequence divergence, and full support from sequencing technologies expected to produce fewer errors of the relevant type. In addition, the strong observed leading/lagging strand bias suggests a genuine biological phenomenon, as does the absence of a strong effect in yeast. Furthermore, a strong hotspot of T→G mutation that has been characterized in the laboratory (Horton et al. 2021) appears to be an example of the phenomenon, as the position is preceded by a run of four G nucleotides.

Selection is expected to affect the estimates of mutation rates. However, its effect is expected to be weak for the NCBI Pathogens data because the mutations analyzed are of recent origin. The ratio of nonsynonymous to synonymous changes in the *Salmonella* is only ~20% lower than expected under selective neutrality, indicating that the effects of purifying selection are modest. Furthermore, only systematic differences in selection between categories of sites will affect the estimates of their relative mutation rates.

### Mechanism

One mechanism of T→G mutation involves incorporation of 8-oxo-G opposite an A in the template. In *E. coli* with inactivated *mutT*, this pathway is the main cause of mutation. In experiments involving such strains, T→G mutation was no more common at sites preceded by G·C runs. This observation is strong evidence that this pathway is not the predominant mechanism of GGGT→GGGG mutation in a wild-type background since it does not disproportionately affect such sites.

Homopolymer runs, particularly of G·C, are prone to expansion and contraction in bacteria. Insertions and deletions in these runs are thought to occur mainly through slipped-strand mispairing (Levinson and Gutman 1987). This process might also be involved in the high rate of T→G transversion after a G·C tract. After slipped-strand mispairing and addition of an extra G at the end of the run, a return to in-register pairing would result in a G:A mismatch and potentially a T→G mutation. Analogous mechanisms would also explain the small effects of preceding G runs on A and C mutation to G. It is not obvious, however, why the effect would be strongest for mutation of T, or why preceding runs of C would not increase the rate of T→C mutation.

The leading/lagging strand asymmetry of the mutation rate suggests that a replication-associated event is involved in the phenomenon It does not, however, distinguish between an event affecting GGGT synthesis that is more common during leading strand synthesis and an event affecting ACCC synthesis that is more common during lagging strand synthesis. Furthermore, orientation might affect a different step in the pathway to mutation from that affected by homopolymer runs.

If G runs do exert their effect during DNA synthesis, they might do so during either replication or repair. Even if slipped-strand mispairing plays no role, it seems more likely that they affect synthesis of the T than that of the complementary A. In the structures of both DNA polymerases 1 (PDB 1QTM; (Li et al. 1999)) and 3 (PDB 3F2B; (Evans et al. 2008)) complexed with primer/template and dNTP, the downstream base(s) of the template strand is/are distant (more than 9 Å) from the template base, incoming dNTP, and polymerase catalytic site. In contrast, the immediately upstream bases of both strands come within 3.5 Å of the nascent base pair. The upstream duplex approximates an ordinary double helix, with interactions that could communicate effects of further upstream bases to the region of the nascent base pair. Thus, the G run seems more likely to affect synthesis when it is upstream of the nascent base pair, i.e., during T-strand synthesis. An effect on T synthesis is also concordant with the fact that the mutation rate is higher when GGGT is on the leading strand, coupled with the suggestion that leading-strand synthesis is more error-prone (Fijalkowska et al. 1998).

### Consequences

The phenomenon accounts for some of the strongest effects of neighboring bases on mutation rate in *Salmonella* (fig. 4). The mutagenic effects of Dcm and Dam methylation explain some, though not all, of the remaining strong effects.

The contribution of the phenomenon to mutation varies among bacteria due to variation in the degree to which mutation rate is elevated, differences in the frequencies of the motifs affected by it, and differences in the base rate of T→G mutation relative to rates of other types of single-base mutation. In *Salmonella*, the phenomenon accounts for 2.42% of mutations, more than half the fraction due to transitions at Dcm-methylated positions. It accounts for more than one third of T→G mutations, and about 10% of mutations from T or A to C or G. The resulting mutations increase the length of the G run by at least one nucleotide, sometimes producing a motif even more prone to mutation and further extension of the run.

Because a highly mutable motif will tend to be short-lived in the absence of selection against the high-frequency mutation, these motifs may be found disproportionately at positions where such selection operates. This is particularly true of sites with more than three preceding G residues, which have particularly high mutation rates. Purifying selection may therefore eliminate most changes at such sites in the long run, though they are apparent in the short-range comparisons considered here.

## Methods

### NCBI Pathogens Data

Most analyses were based on data from the NCBI Pathogens database (https://www.ncbi.nlm.nih.gov/pathogens/). This contains information derived from whole-genome sequencing of large numbers of bacterial isolates for many taxonomic groups (species or genera; *Shigella* is combined with *E. coli*, of which it is part phylogenetically). It provides single-nucleotide polymorphism (SNP) calls for clusters of very closely related isolates, along with phylogenetic trees based on these SNPs.

The build runs (versions of taxon-specific datasets) used in the analysis were PDG000000002.2479 (*Salmonella*), PDG000000004.3383 (*E. coli*), PDG000000001.2875 (*Listeria*), PDG000000003.1716 (*Campylobacter*), PDG000000032.298 (*Neisseria*), PDG000000036.681 (*Pseudomonas aeruginosa*), PDG000000055.317 (*Vibrio cholerae*), PDG000000026.169 (*Legionella pneumophila*), and PDG000000073.309 (*Staphylococcus aureus*).

The sequencing technology used for each assembly was determined from the “Sequencing technology” line of the associated assembly_stats.txt file. For mutations that were inferred to have occurred on an internal branch of the tree, it was checked that the SNP data for the supporting genome(s) sequenced by the technology of interest contained the derived G, rather than an ambiguity that had been resolved by the ancestral state reconstruction or a different nucleotide due to an ostensible second change at the position.

### Reconstruction of Sequence Evolution

For each cluster containing at least five isolates, sequence changes within the cluster were reconstructed by maximum parsimony. The nature of each change (the ancestral and derived nucleotide) and its sequence context were recorded, along with its location in the reference genome for the cluster The details of this procedure have been described elsewhere (Cherry 2018; Cherry 2020).

### Analysis of Apparent Mutation Rates

Sequence changes that were mapped to branches descending directly from the root of a tree were excluded from the analysis because the impossibility of determining their direction (e.g., C→T vs. T→C). Some were also excluded on the basis of information about sequence reads that aligned to the SNP position. For cases in which more than one isolate was affected, a single representative was chosen at random so that all mutations could be subjected to identical criteria. Only cases in which there were at least 20 aligned reads, at least 90% of them supported the called base, and at least 25% were supporting reads in each direction were included in the analysis.

Numbers of T→G changes at positions with various numbers of preceding G residues were calculated, as were analogous counts for other types of mutation. For analyses involving multiple NCBI pathogens clusters, frequencies of sites of each type were computed as averages among the reference genomes for each cluster, weighted by the number of usable changes in the cluster. The weighted mean site frequencies were multiplied by the weighted genome size. This multiplication does not affect relative rates at different types of sites, since it affects all sites in the same way, but it allows interpretation of the products as approximate numbers of sites per genome. Simply using the site counts in a single genome yielded similar results, so the details of this procedure are unimportant. Numbers proportional to rates per site were calculated as the ratio of relevant changes to the (weighted mean) number of occurrences of the site category. Rates relative to positions adjacent to zero-length tracts were calculated by simple division. The relative rate for zero-length tracts is identically one, and hence not presented, except in the analysis of leading/lagging strand asymmetry, for which the denominator was calculated using both strands.

### Origin and Terminus of Replication

The approximate locations of the origin and terminus of replication for CP093400.1 (PDT001274653.1), the reference sequence for *Salmonella* cluster PDS000089910.229, were determined from GT skew. This was calculated for each position in the chromosome as the count of G and T nucleotides minus the count of C and A nucleotides in the sequence up to that position. The positions with minimum and maximum GT skew were taken as the origin and terminus respectively.

### Analysis of *E. coli* Laboratory Evolution and Mutation Accumulation Data

Mutation data from Couce et al. (2017) was downloaded from https://datadryad.org/stash/dataset/doi:10.5061/dryad.sq67g. It was analyzed in conjunction with the applicable chromosome sequences of *E. coli* REL606 (NC_012967.1) and MG1655 (NC_000913.2). Sequence differences appearing in multiple clones from the same experimental replicate were counted only once for the purpose of estimating relative mutation rates. For assessing the relationship between changes of interest and time, every difference from the REL606 sequence in every genome from a time point was counted.

### Analysis of *Pseudomonas aeruginosa* Serial Isolate Data

The data provided by Marvig et al. (2015) was used for the analysis. Isolates bearing nonsynonymous changes or insertions or deletions in mutator genes were considered to have a mutator phenotype. Point mutations present only in such isolates were excluded from the analysis.

The isolates are divided into 36 clone types. Because a single mutation event may affect more than one isolate from a clone type, counts from isolates within a clone type are not independent. Therefore, just one non-mutator isolate was chosen randomly from each clone type for the purpose of calculating the coefficient of correlation between the number of mutations of interest and the number of point mutations of other types. Correlation coefficients were calculated for 100,000 independent random choices of isolates.

### Randomized Counts for Correlation Coefficients

Correlation coefficients between the numbers of mutations of interest and of other types of mutation were calculated for *Salmonella* and *Pseudomonas aeruginosa*. Because the actual numbers of mutations vary stochastically around their expected values, the correlations between the counts will usually be imperfect even if their expected values are exactly proportional. For comparison, the correlation coefficient expected under such proportionality was estimated using each randomly chosen set of isolates or isolate pairs. For each set, counts were drawn from a multivariate hypergeometric distribution such that the number of mutations for each isolate (pair) and the total number of mutations of interest were maintained. This procedure is equivalent to randomly designating mutations affecting the chosen isolates as mutations of interest, with the total number of mutations of interest equal to the actual total and each mutation equally likely to be so designated.

## Acknowledgments

This work was supported by the intramural research program of the National Library of Medicine, National Institutes of Health. The opinions expressed in this article are those of the author and do not reflect the view of the National Institutes of Health, the Department of Health and Human Services, or the United States government.

